# In-Silico Stability Predictors: Investigation of Performance Towards balanced Experimental Data

**DOI:** 10.1101/2025.03.28.645695

**Authors:** Kristine Degn, Mattia Utichi, Pablo Sánchez-Izquierdo Besora, Matteo Tiberti, Elena Papaleo

**Affiliations:** Cancer Systems Biology, Section for Bioinformatics, Department of Health and Technology, Technical University of Denmark, 2800, Lyngby, Denmark; Cancer Structural Biology, Danish Cancer Institute, 2100, Copenhagen, Denmark

## Abstract

Change in protein stability, quantified as the change in Gibbs free energy of folding (ΔΔG) in kcal/mol, plays a crucial role in functional alterations of proteins, with misfolding and destabilization commonly associated with pathogenicity. The past two decades have brought the development of bioinformatics tools leveraging evolutionary knowledge, empirical force fields, and machine learning to predict stability alterations. However, existing tools are often optimized towards or trained on limited experimental data, leading to unbalanced datasets and potential overfitting. The research objective is to benchmark selected stability predictors using an unbiased and balanced dataset, with AlphaFold structures as input. We demonstrate a performance decline when balancing data across amino acids, stabilizing and destabilizing mutations, and protein representatives, highlighting that redundancy alone is insufficient for benchmarking correction. Additionally, we illustrate that a protein structure ensemble from molecular dynamics acts as a superior input compared to a single static structure. At the same time, coarse-grained methodologies tend to decrease the output quality.

## Introduction

The structural stability of a protein is a fine-tuned equilibrium allowing dynamic interaction while retaining functional integrity^1,2^. Hence, destabilizing amino acid substitutions may disrupt the stability of the protein, yielding the protein defective or otherwise functionally altered^3,4^. Such alterations might be associated with pathogenicity^5–7^. Accordingly, experimental approaches to assess substitution-based stability alterations, measured as change in folding Gibbs free energy between mutant and wild-type, ΔΔG, have been developed^2,8–10^, along with computational predictors of ΔΔG utilizing the protein sequence^11–16^ or relying on the three-dimensional structure of the protein^16–34^.

Structural bioinformatics is currently experiencing a leap forward^35^, utilizing sophisticated machine learning methodologies to describe protein alterations in greater detail^21,36,37^. Despite this progress, the performance of stability predictors upon mutation seems to have stagnated and under-performs when encountering previously unseen data generated after model training or not included in tool optimization^38^. Tools rooted in biophysics, such as Rosetta^17–19^ and FoldX^20^, have outperformed the machine learning models when tested on such independent datasets^38^, likely because the machine learning methodology has been trained on unbalanced datasets, indicating that the available data was not large and diverse enough for data-driven feature selection^39^. Performing experiments that measure structural stability is expensive and time-consuming, and even when collected, the data may not be shared or formatted in a reusable way^40^. The currently available datasets were compiled from various independent sources in the literature^41^, with each having a different scope and scientific purpose. Consequently, in such datasets, there are typically only a few mutations occurring on any given protein, and they might not be diverse enough in terms of representing protein families and mutation types^41^. This results in both data bias and non-uniformity. For these reasons, the publicly available data pool has several limitations^42^. For instance, the experimental data sets include experiments performed using diverse protocols and experimental conditions^27,42^. They show a substantial bias toward certain types of mutations, i.e., substitutions to alanine^27,42^, and are biased towards destabilizing mutations^42,43^. Additionally, the mutation distributions across the screened proteins are non-uniform, where a small group of proteins accounts for most of the datasets^38^.

Computational approaches have historically introduced methodological bias during development, such as neglecting anti-symmetry^41–43^, yielding cases in which a mutation and its reversion to wild-type have significantly different absolute changes of stability. It has been argued that the lack of anti-symmetry is an artifact of the poor estimation of stabilizing mutations^43^. An additional source of computational bias, leading to an inflated performance estimation for the predictors, is sequence or structural similarity between proteins used in training and test datasets^42,44^. Indeed, this challenges classic cross-validation, as the developer may need to filter proteins or mutations to accurately measure performance^38^. Most of the experimental data used to train, optimize, or test the different tools has traditionally been collected in the Protherm database, risking overlap of training and testing data between methods^38,40,45^. A turning point in the availability of experimental data occurred in 2023 when Tsuboyama et. al.^2^ introduced a novel technique allowing high-throughput free energy of folding (ΔG) estimation on an unprecedented scale. They obtained the measurements on small model systems, specifically proteins 60-80 amino acids in length, without cofactors or ligands. Since then, computational methodologies leveraging this data to predict ΔG and ΔΔG’s have emerged^46–48^, allowing researchers to apply tools developed for this new data to prior datasets to investigate the transferability of performance.

Applying stability predictors to study the mutational impact on function has a wealth of purposes, such as variant effect prediction and classification^49^. Given the nature of stability’s relation to protein function, evaluating the computational methodologies against experimental data through a benchmarking study can aid researchers in choosing which tool to apply to their proteins. Unbiased benchmarking, therefore, provides a solid foundation to assess the quality of the tools in the current context. Stability prediction benchmarking has been conducted continuously over the years^44,50^, and the performance landscape has indeed changed since the release of the high-throughput data^2^. Nonetheless, we would like to address three factors that, we believe, have so far been overlooked: i) effect of the source input structure on predictions. While researchers regularly use predicted models^36,49,51,52^, most benchmarking is conducted on experimentally determined protein structures^50,53,54^. In addition, case studies have illustrated that ensembles of structures obtained from molecular dynamics simulations yield better outcomes when used as input than single structures^55,56^. We aim to evaluate performance on predicted structures generated by AlphaFold2 and examine how the molecular dynamics ensemble derived from the AlphaFold2 model influence this performance. Additionally, we investigate whether this performance improvement extends to ensembles obtained with coarse-grained methodologies, specifically CABS-flex^57^, which introduces random movements of the backbone to emulate movements of a structure solved using NMR. ii) Amino Acid Balance. Most datasets released do not address the data imbalance caused by alanine scans and effectively risk benchmarking the predictors towards their ability to represent mutations to alanine. iii) Protein Balance. Along the same lines as the amino acid balance, our dataset should be equally representative of as many protein folds as possible and not be over-represented with mutations from a few specific proteins, which might be available for historical reasons. Thus, here we investigate and benchmark the change in stability predictions from Rosetta^17–19^, FoldX^20^, RaSP^21^, DDgun seq^11^, DDgun 3D^11^, and ThermoMPNN^47^. We benchmark on an unbiased dataset derived from Pancotti et al.^54^, using AlphaFold2^37^ structures as input.

## Results

### Performance of Predictors is inflated on unbalanced Data

Zheng et al.^50^ introduced the S4038 benchmarking dataset in 2023, derived from ThermoMutDB^58^, FireProtDB^59^, and ProThermDB^60^, and curated to exclude overlap with the widely used S2648 dataset^61^. They evaluated 27 predictors across five performance metrics, concluding that ThermoMPNN^47^ demonstrates the most consistently good overall performance. They also highlight significant performance biases, noting that an imbalance between stabilizing and destabilizing mutations in training, testing and benchmarking may lead to inflated performances. ThermoNet^30^ performs best when predicting stabilizing mutations, although its accuracy remains lower than when predicting destabilizing mutations. While S4038 addresses redundancy, it does not mitigate biases such as amino acid overrepresentation (**Figure 1A**), class imbalance (**Figure 1B**), or reliance on specific proteins (**Figure 1C**).

**Figure 1:**
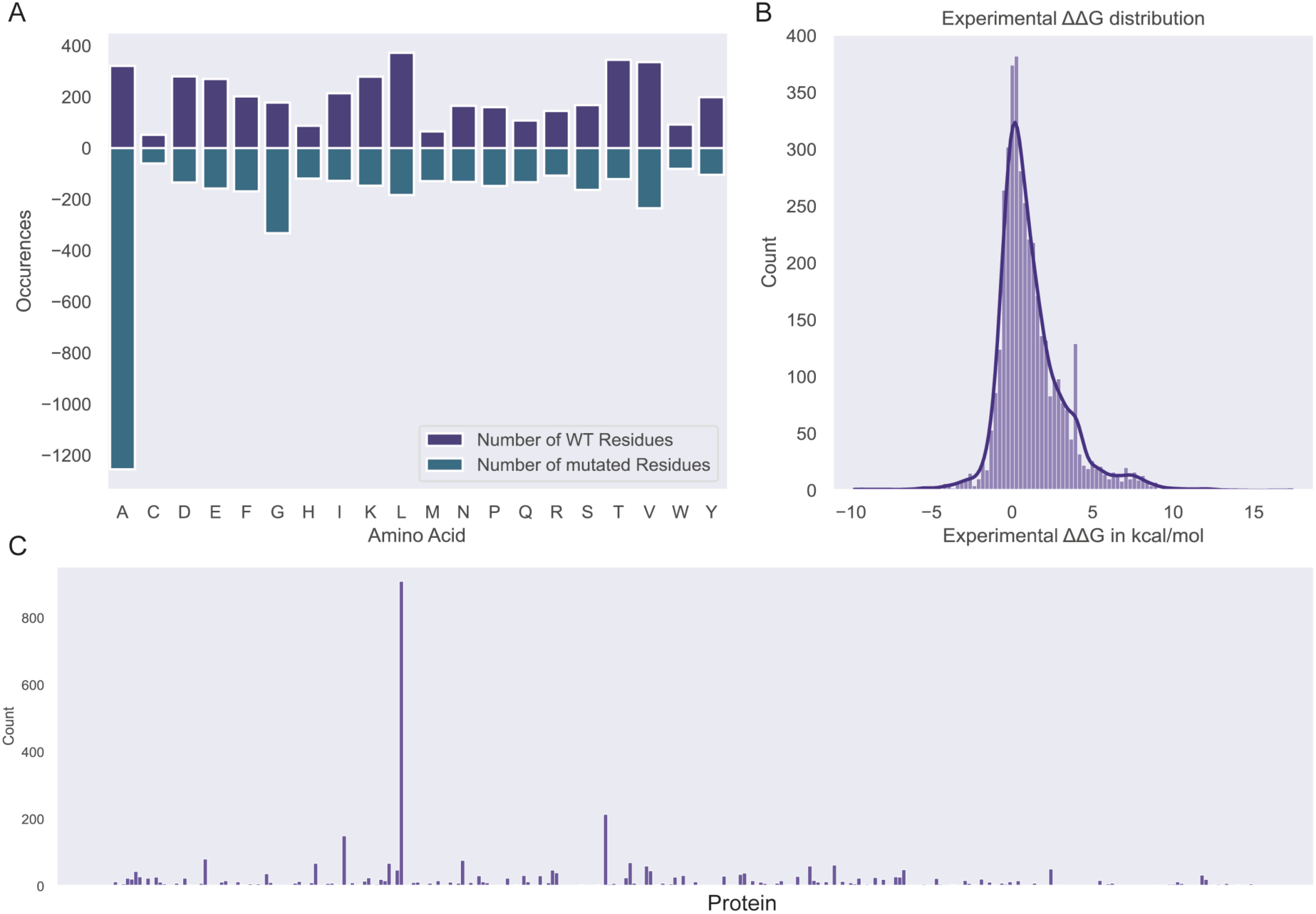
Distributions of S4038 (A) Amino Acid count. The plot illustrates the number of sites mutated from a wild- type (WT) amino acid on residues of the sites in the dataset in purple, positive y-axis, and number of sites mutated to an amino acid, in teal, negative y-axis. Over 1200 out of the 4038 mutations are mutations to alanine, potentially benchmarking towards alanine scans. (B) The histogram illustrates the distribution of the experimental ΔΔG values, showing a higher number of destabilizing mutations (ΔΔG > 0) than stabilizing mutations (ΔΔG < 0) (C) The bar chart illustrates each of the proteins in the S4038 dataset on the x-axis and the number of mutations reported in each protein. It is evident that one protein alone accounts for > 800 of the mutations.

The authors propose expanding the dataset to increase the representation of stabilizing mutations^50^. We aim to tackle this issue as an overrepresentation and employ biased under- sampling to rebalance S4038, achieving a more uniform distribution of mutations across proteins, amino acid substitutions, and stabilizing versus destabilizing mutations. While this reduces dataset size, potentially affecting statistical significance and reliability, we mitigate this by performing ten rounds of this under-sampling to produce ten balanced subsets. As this does not overcome the challenge of having fewer data points, thus potentially lowering the reliability of the results, we also generate ten random subsets of comparable size as controls to isolate any effects due to reduced data size. Pearson Correlation Coefficients (PCC) between experimental data and predictive methods reveal decreased performance for all predictors across S4038, the balanced subsets, and the random subsets (**Figure 2**), indicating that the lowered performance of the balanced subsets is indeed not an artifact of the lower number of data points, but rather a suggestion of overfitting or overreliance on unbalanced data.

**Figure 2:**
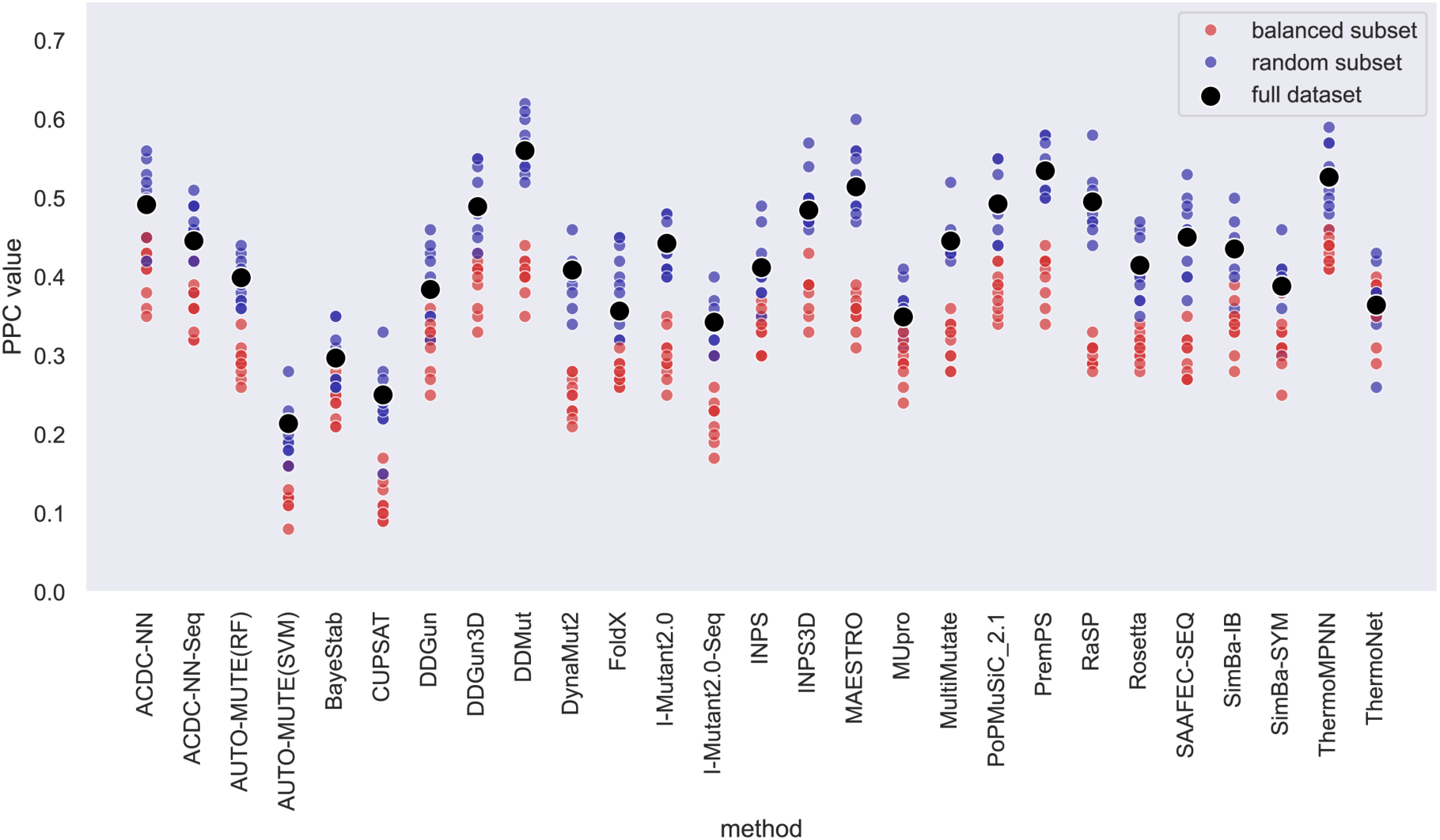
A scatterplot of the Pearson Correlation Coefficient (PCC) of the full S4038 dataset (black), and the balanced subsets (red) and random subsets (blue). For many of the benchmarked tools (x-axis), there is a clear separation between the balanced subsets and the random subsets, where the balanced subsets tend to have lower PCCs towards the experimental data.

Overall, Pearson Correlation Coefficients (PCC) tend to be higher for random subsets than for balanced ones, with some predictors affected more than others. For instance, RaSP^21^ and DDMut^24^ show markedly lower performance on balanced subsets, while ThermoNet^30^ maintains similar performance across both, consistent with its strong performance on stabilizing data, as reported previously^50^. ThermoMPNN^47^ outperforms other predictors on balanced data from Zheng et al.^50^, with an average PCC of 0.43, followed by PremPS^29^ and DDMut^24^ at 0.40. Using a one-sample t-test, we compare the PCC of the full dataset with that of both random and balanced subsets. Results show no statistically significant difference between the full dataset and random subsets (p > 0.05), but significant differences with balanced subsets for all tools except ThermoNet (p = 0.85). An independent t-test between random and balanced subsets confirms significant differences across all tools except ThermoNet (p = 0.96). These results suggest that dataset balancing produces notably different performance outcomes to the full dataset, revealing potential inflation in stability predictor benchmarks. Although S4038 is constructed to avoid overlap with the S2648 dataset, some overlap with training sets persists. Aware of this, Zheng et. al annotated the data points, and after excluding all training-related mutations, 568 unique mutations remained (**Figure S1**). Rankings remain consistent, with ThermoMPNN leading (PCC = 0.34), followed by INPS3D (0.33) and DDMut (0.31). When rebalanced, the balanced and random subsets show no significant performance separation, indicating that prior differences may primarily stem from overfitting. On balanced, unbiased data, AUTO-MUTE(RF)^33^ achieves the highest PCC (0.31), followed closely by ThermoMPNN (0.30), with minimal performance reduction due to balancing.

### Benchmarking Predictive Performance with AlphaFold Structures

In 2022, Pancotti et al.^54^ curated a benchmarking dataset of proteins from the ThermomutDB^58^, addressing the redundancy caused by using similar proteins for both model development and testing, which leads to overestimating the performance of the predictors. They extracted variants from the ThermoMutDB, with less than 25% sequence similarity to well-known datasets^34,62^, and manually cleaned the data to exclude inconsistencies, providing a dataset named S669. We further filtered the dataset to include only proteins with available AlphaFold models in the AlphaFold database and mutations located within domains with a pLDDT score > 70 (see Methods for details). This filtering resulted in a final dataset of 612 mutations, referred to as S612 in this work. We then applied FoldX in the Mutatex implementation^56^, Rosetta cartesian in the RosettaDDGpredict implementation^63^, RaSP^21^, DDGun^11^, DDgun3d^11^, and ThermoMPNN^47^ to the S612 dataset (**figure 3**, **Table 1**).

**Figure 3:**
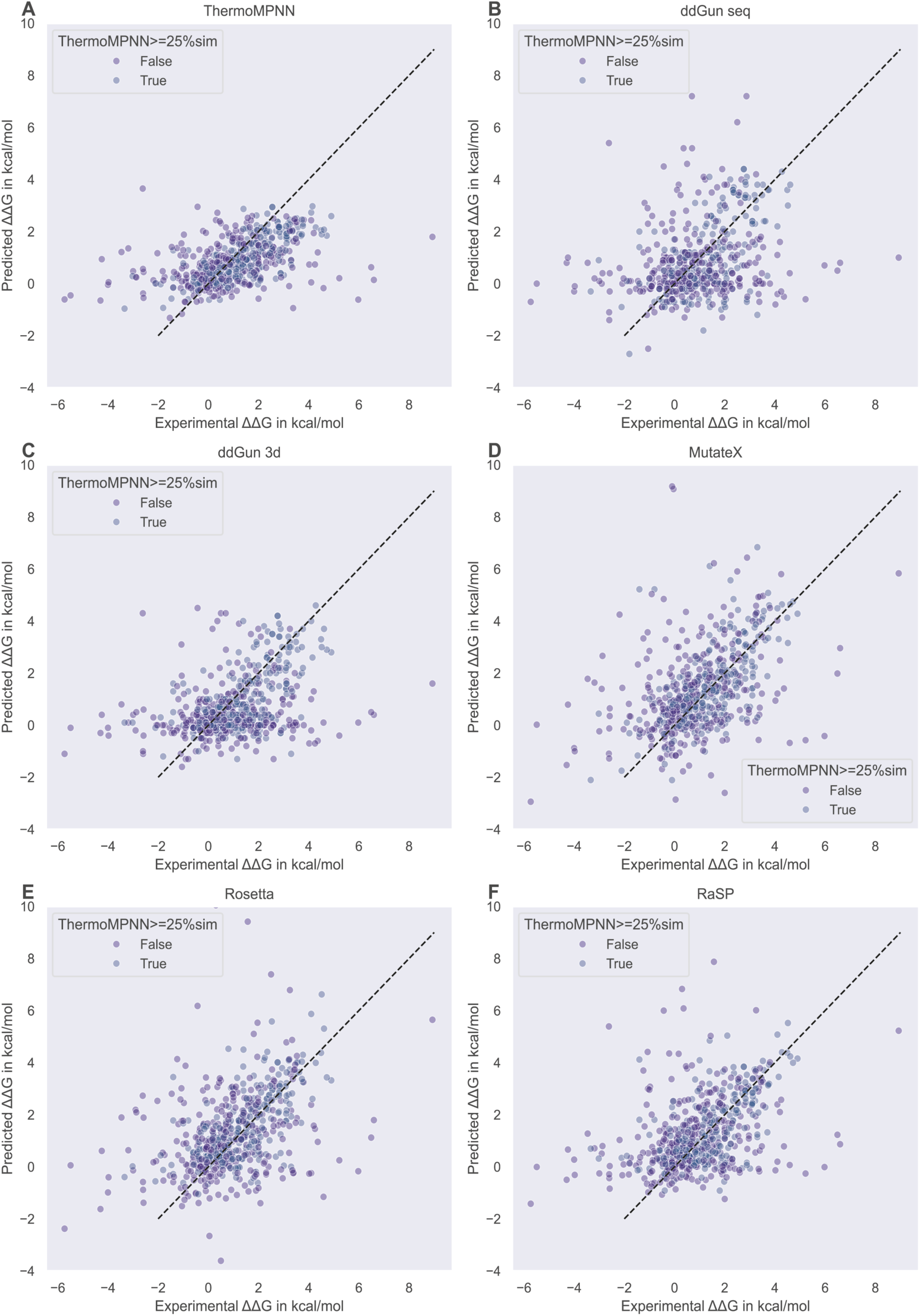
All scatterplots display the experimentally determined change in stability on the x-axis and the predicted values of each tool on the y-axis. The black like indicates perfect correlation. Blue dots indicate data points with a 25% or higher sequence identity to the training set used in ThermoMPNN.

**Table 1.**

|  | S612 |  |  | 332 mutations (< 25% sequence similarity to proteins used to train ThermoMPNN) |  |  |
| --- | --- | --- | --- | --- | --- | --- |
|  | PCC | SCC | MAE | PCC | SCC | MAE |
| Mutatex | 0.44 | 0.50 | 1.20 | 0.33 | 0.37 | 1.50 |
| Rosetta | 0.44 | 0.47 | 1.17 | 0.31 | 0.34 | 1.46 |
| RaSP | 0.42 | 0.45 | 1.14 | 0.30 | 0.34 | 1.40 |
| DDGun Seq | 0.29 | 0.31 | 1.31 | 0.13 | 0.16 | 0.15 |
| DDgun 3d | 0.38 | 0.37 | 1.16 | 0.16 | 0.20 | 1.38 |
| ThermoMPNN | 0.51 | 0.58 | 0.96 | 0.36 | 0.44 | 1.15 |

In the original paper on ThermoMPNN^47^, they report a PCC of 0.43 for their performance towards S669, indicating that the version of ThermoMPNN they benchmarked towards is a version trained with a homolog-free megascale training dataset to avoid data leakage and that S669 is filtered down to 420 variants to avoid overlap to the training data. We also filtered the dataset based on the ThermoMPNN’s training data to only include proteins with less than 25% sequence similarity. The filtering step results in fewer mutations (332 mutations) due to two key factors: first, we obtained ThermoMPNN results from the published version of the tool, thus not with leakage-free training, and second, we started with 612 mutations compared to the full 669 mutations. The correlation decreases for this filtered data on all the predictors. We observe a Pearson correlation coefficient of 0.51 overall and 0.36 when removing structures comparable to the training data. 0.36 is a similar performance to the 0.34 found for the unbiased data of the S4038 dataset. The S612 and S4038 datasets share 57 proteins, while 20 are unique to S612 and 46 to S4038. All mutations and their predicted values are available in the supplementary materials as tableS1.

Benchmarking these tools against the dataset reveals that ThermoMPNN outperforms the energy-based tools. Additionally, RaSP shows a correlation comparable to Rosetta and FoldX in the MutateX implementation while achieving a lower MAE. In 2023, Gerasimavicus et al.^64^ found that ‘foldetta’, an average of FoldX and Rosetta, outperformed each of the tools individually. Our results align with this finding: ‘foldetta’ achieves a Pearson correlation coefficient of 0.48 in S612 and 0.35 in the filtered dataset, both better than the individual tools. This is also true when substituting Rosetta with RaSP, where we find correlations for the ‘foldaSP’ at 0.47 and 0.34 respectively. This indicates that each tool individually has some limitations that are mitigated through combination approaches, hence the field of stability predictions may benefit from ensemble methodologies, as seen in other variant effect predictor methodologies, such as REVEL^65^. As with S4038, these correlations are dependent on the input data. Despite efforts to have very limited redundancy, other imbalances persist, such as an overrepresentation of alanine scans and a majority of destabilizing mutations (**figure 4C, 4A**). To address these issues, we balanced the S612 dataset mirroring the balancing of the S4038 dataset (**figure 4E**) and its filtered version with 332 mutations (**figure 4F**). For the S612 dataset, we find a less pronounced separation of the balanced and random subsets, yet still find a better performance in the random datasets. In contrast, the filtered version with 332 mutations showed no indication of bias. The top five performing methods are all combination methods, with the average of MutateX and ThermoMPNN (PCC=0.37), MutateX, RaSP and ThermoMPNN (PCC=0.37), MutateX, Rosetta and ThermoMPNN (PCC=0.37), MutateX, Rosetta, RaSP and ThermoMPNN (PCC=0.36), and MutateX, Rosetta, Ddgun3D and ThermoMPNN (PCC=0.36).

**Figure 4.**
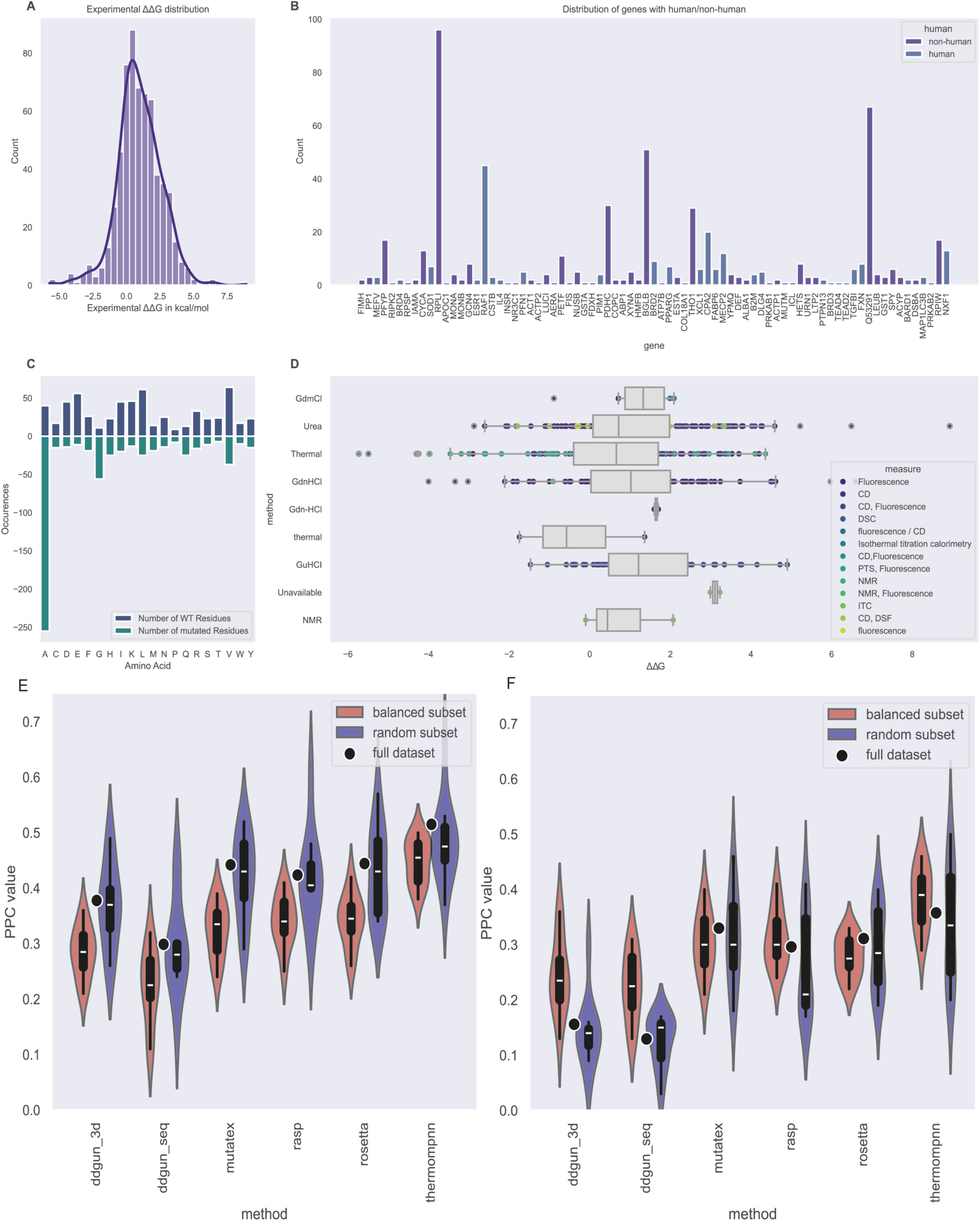
(below): The distribution of the S612 dataset. (A) A histogram showing the distribution of the experimental change in stability kcal/mol. We see an overweight of destabilizing mutations, i.e., ΔΔG > 0. (B) A barplot of the proteins in the dataset. We see that some proteins are more represented than others, relying on a few proteins for most of the dataset. (C) A barplot of the substitutions, where every amino acid is represented on the x-axis, and the number of wild-type residues in the datset of that amino acid is represented on the positive side of the y-axis in teal. The substituted aminoacid is represented on the negative part of the y-axis illustrating a clear overreliance on alanine. (D) A boxplot illustrating the experimental methods and measure used to obtain the data to investigate if specific methodologies tend to over or underestimate. There is no evidence that the methodologies bias the data. (E) A violinplot of the Pearson Correlation Coefficient (PCC) of the full S612 dataset (black), and the balanced subsets (red) and random subsets (blue). There is less of a separation between the balanced subsets and the random subsets than we find in S4038. (F) for the filtered 332 mutations, the violinplot indicates no advantage of the random subsets compared to the balanced subsets.

### Assessment of the Performance of Ensembles as Input with CPA2 as a model system

The CPA2 gene is translated into Carboxypeptidase A2, a secreted catalytic enzyme. CPA2 contains two domains connected by a short linker. The first domain (residues 23-94) has been studied as a model system, as it is a folded domain without any catalytic or binding sites. In the S612 data, 20 mutations originate in CPA2, all in this short domain (Sites indicated in **figure 5A**). Additionally, experimental data on this domain, published on the MaveDB^66–68^, from the mega-scale dataset^2^, measure variant effects on folding. We calculated ΔΔG values by comparing them against the wild-type value. Correlation with the ΔΔG values from the S612 dataset yielded a Pearson correlation coefficient of 0.80 (**figure 5B**), suggesting that the two datasets are in strong agreement. Among the human proteins in the S612 dataset, only CPA2 and ATP7B have associated depositions in the MAVEdb. Since ATP7B only covers a single mutation from the S612 dataset, we chose to investigate CPA2. The ΔΔG scores from the mega-scale dataset^2^ show a bimodal distribution (**figure 5C**), with one peak around the wild-type value and another near 3.8 kcal/mol, representing destabilizing variants. However, when double mutations, insertions, and deletions are removed, leaving only the single mutations studied here, the distribution shifts to center around the wild-type value, with a tail towards the destabilizing variants (**figure 5D**).

**Figure 5:**
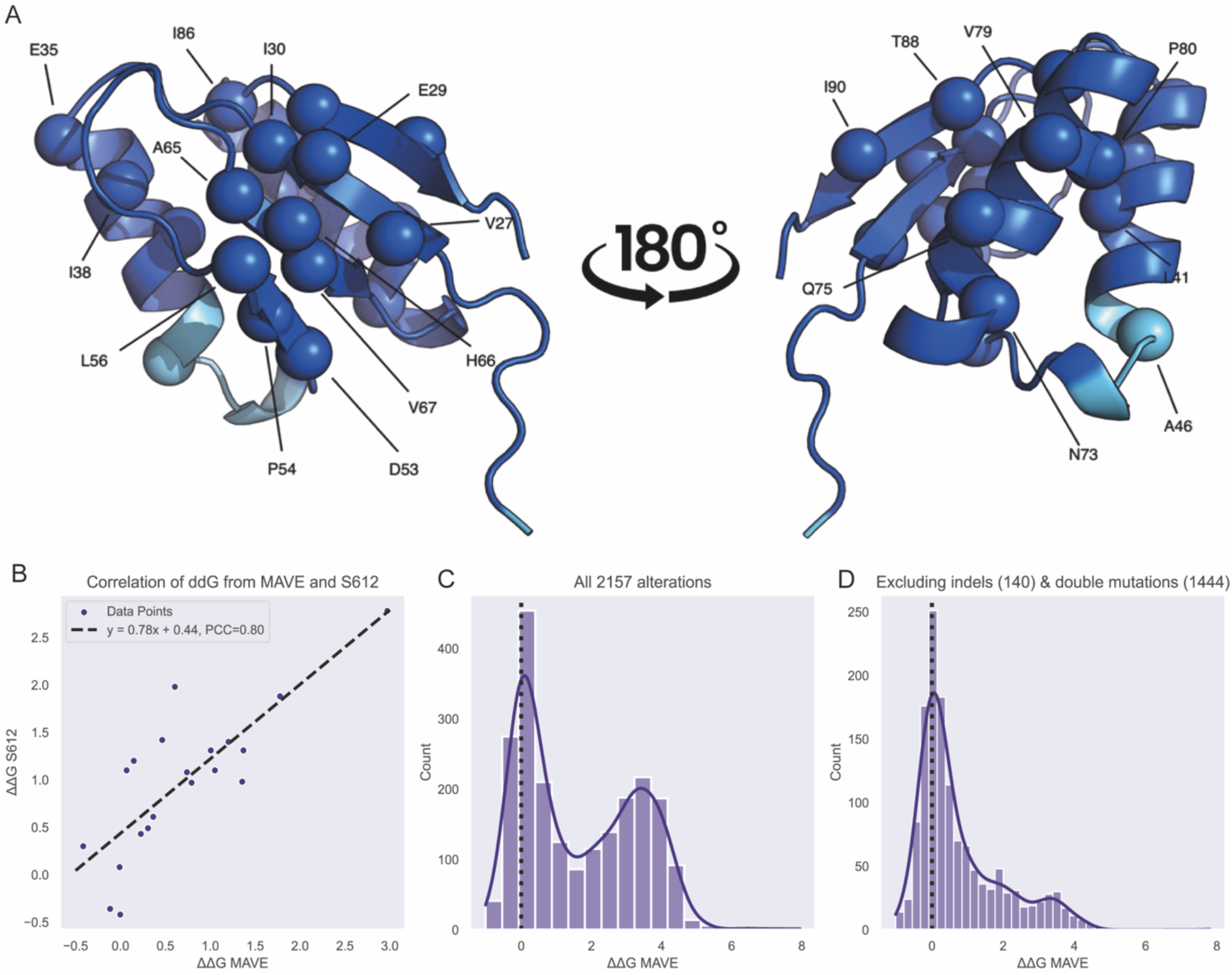
Overview of CPA2 data (A) is the CPA2 domain (residues 23-94) where each mutated site from the S612 dataset is annotated. (B) The correlation between the ΔΔG (kcal/mol) from the mega-scale dataset^2^ on the x-axis and the value reported in the S612 dataset on the y-axis for the 20 mutations. We see a correlation between the two. (C) the ΔΔG values of the mega-scale data where 0 kcal/mol indicates no change, a positive value indicates a change in stability of a variant causing a destabilizing effect and a negative value a stabilizing effect. We see a peak around 3.8. (D) Once removing insertions, deletions and double mutations, the number of destabilizing mutations is reduced, and only single mutations are retained.

To investigate the impact of starting structure on the calculations, we performed saturation mutagenesis of this domain with MutateX, RaSP, and ThermoMPNN, on a single AlphaFold model and on two ensembles of structures: one generated with Molecular Dynamics and one with the coarse grain method CABSflex^57,69^. We have previously shown how introducing a molecular dynamics ensemble of p53 improved FoldX results^55^, as this overcomes the limitation of the fixed backbone. For CPA2, we also see an increase in the Pearson correlation coefficient between the simple (single structure) and MD ensemble (**figure 6**). Interestingly, when introducing mutations, the coarse grain models generated by CABS-flex yield a worse Pearson Correlation coefficient than the single AlphaFold model. This suggests that there is seemingly no performance increase by random moves of the backbone. This pattern is mirrored with both RaSP and ThermoMPNN, yet none improves as much as FoldX. In particular, RaSP only increases the Person Correlation coefficient by 0.02 points, while the improvement is 0.07 points for FoldX, and 0.06 points for ThermoMPNN. As CPA2 is part of the mega-scale dataset and can be part of the data on which ThermoMPNN is trained, the correlation itself is likely inflated. The correlations for the 20 mutations from the S612 dataset can be found in the supplementary materials and follow the same pattern (**figure S2**). Notably, averages of multiple methods do not outperform either RaSP or ThermoMPNN.

**Figure 6:**
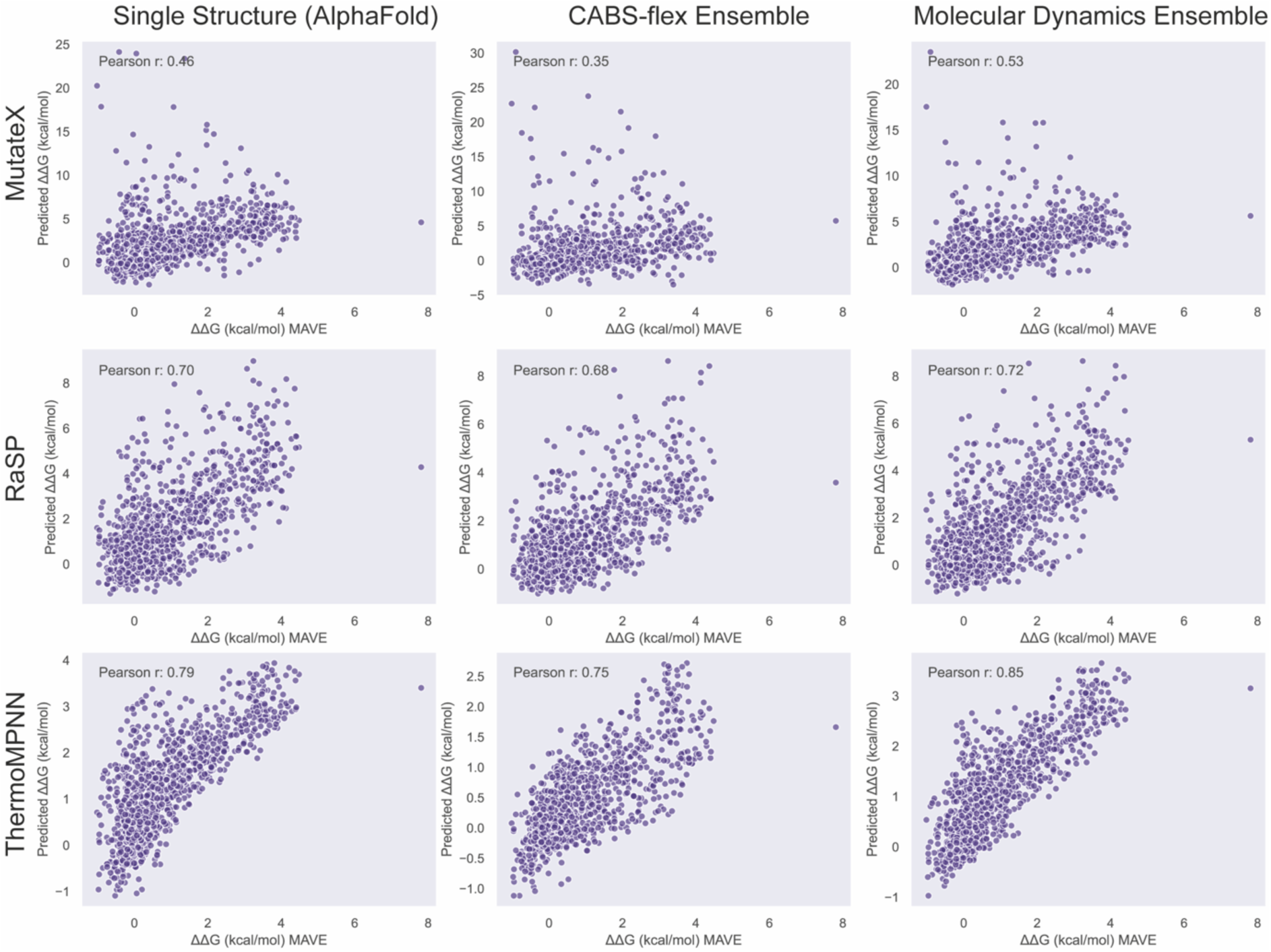
Scatterplots of the ΔΔG values from the mega-scale dataset and the predicted ΔΔG values on three different inputs, the simple (static) structure from AlphaFold, a CABSflex ensemble and a Molecular Dynamics ensemble.

**Figure 7.**
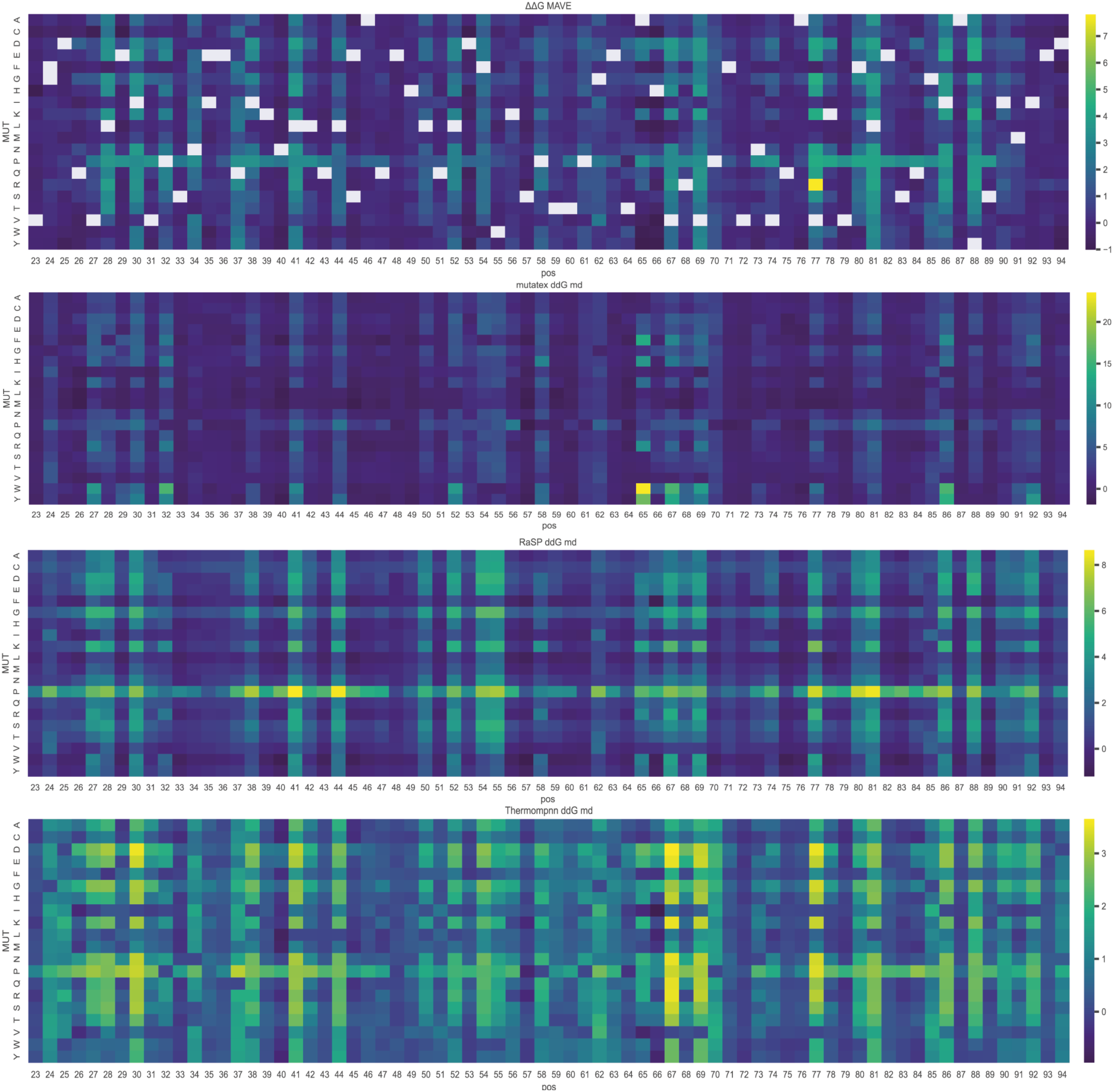
Heatmaps illustrating the change in stability of every mutation in the domain. The first panel is the experimental ΔΔG from the mega-scale dataset, and the following panels are predictive ΔΔG with the MD ensemble as input. We see that all the methods can identify sites where mutations tend to destabilize the protein, as well as the effect of proline. Notice that the scales of the different methodologies are quite different, while only a single datapoint in the mega-scale dataset reach > 7 kcal/mol, the MutateX predictions reach above 20 kcal/mol, while RaSP and ThermoMPNN tend to stay in a lower range, making the smaller differences clearer in the plotting.

Apart from a direct correlation with the experimental data, we also review specific sites or mutations that tend to follow the experimental values. As illustrated in **figure 9**, we see sites in the ΔΔG from the mega-scale dataset, where almost any mutation is destabilizing, such as residues L28, I30, N34, Q37, L41, L44, L52, F54, V67, V69, V77, L81, I86, and Y88, which are all in the domain’s core, with rotameric state towards the center. We also see a clear disfavor of Proline as a substitution. The most notable mutation is V77R, which is a mutation from a small to a much bulkier residue in the narrow space of the core of the domain. This results in a very high value, which appears as an outlier (**figure 8**). Interestingly, none of the computational methodologies detect this mutation. However, they tend to identify the sites of interest and correctly classify core mutations as destabilizing.

**Figure 8:**
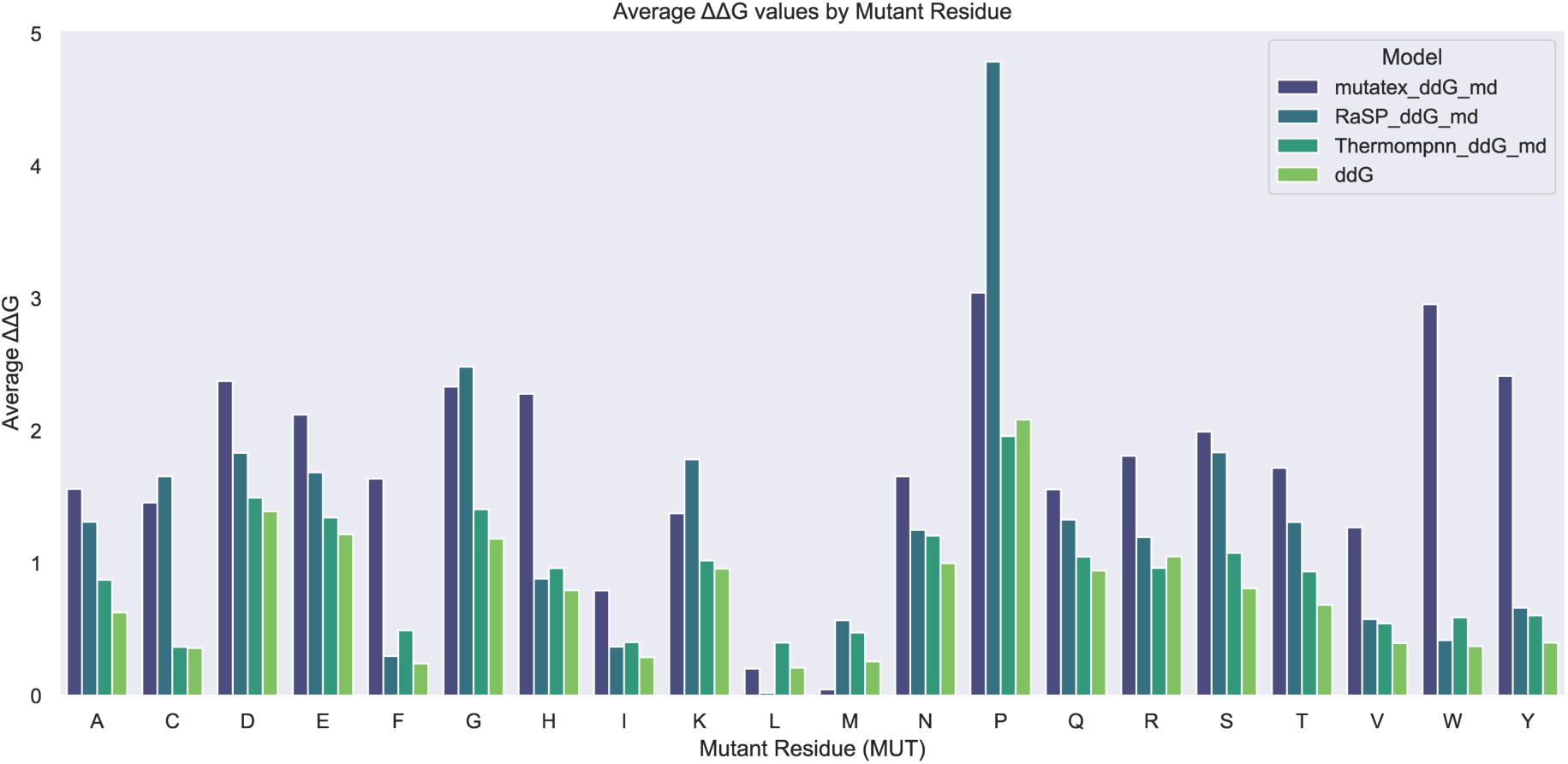
A barplot representing the average ΔΔG in kcal/mol each method assigns an amino acid substitution. The mega-scale experimental value is the light green bar. We see that ThermoMPNN and the experimental data lagely seem to follow each other. Interestingly, we see that RaSP seems to vastly overestimate the effects of Glycine and Proline substitutions. These effects are likely some of the limitations of Rosetta that are amplified rather than negated in the model. While FoldX in general overestimates the effects, this is especially true for Phenylalanine, Tryptophan, Tyrosine, and Histidine which are all amino acids with large side chains, which may be explained by the stiffness of the backbone, even when calculating the average over an MD ensemble.

RaSP disfavors Proline. On average, RaSP appears to estimate Proline as much more destabilizing than the other methodologies (**figure 8**). This could indicate a bias towards Proline and should be considered when evaluating RaSP results. RaSP seems to estimate mutations to Glycine to be more destabilizing than expected, which is expected since it is trained on Rosetta data. ThermoMPNN scale is much closer to the experimental scale, while FoldX estimates values all the way to > 20 kcal/mol. Normalizations for these methods can be found in the supplementary material.

## Discussion

The findings of this study emphasize the significant impact novel data can have on reshaping the current landscape of methodologies, particularly in overcoming challenges that have persisted for many years in their benchmarking. However, it is notable that even with the new mega-scale data^2^, the ThermoMPNN^47^ developers found it necessary to combine the new data with older database data to achieve a good performance with respect to experimental data. This need indicates the persistent requirement for more experimental findings. However, the present findings might suggest some shortcomings of either the tools’ linear combinations or the choice of features (Blossom substitution Matrix^70^, Skolnick’s statistical potential^71^, and Kyte-Doolittle’s scale^72^). We show how FoldX outperforms Rosetta on the S612 dataset. We also observe that RaSP, the tool trained on Rosetta predictions, almost reaches the performance of Rosetta’s predictions, promising findings in terms of computational efficiency as both are less computationally demanding methodologies. We find that predicting the stability change on proteins with sequences having less than 25% sequence similarity to the data used to train or optimize the model provides a fair benchmarking approach. The correlations we observe should be considered as a minimum performance benchmark, meaning that users should not be discouraged by low correlations.

Although the balancing may illustrate some biases, change in stability upon mutation largely indicates inherent destabilization, as evolution has fine-tuned the equilibrium between function and stability – why most changes are likely to be deleterious^73^. Therefore, balancing on this parameter is relevant to evaluate the utility of the tools although it does not necessarily favor the performance. Continuing to assess the predictors on destabilizing mutations and alanine substitutions falsely inflates performance metrics and might lead to over-reliance on results obtained with sub-par predictors. Indeed, we show how the method ranking changes after balancing the S4038 dataset^50^. In comparison to the findings of Zheng et al.^50^, our correlations to experimental data are significantly lower. When we extract their annotations of sequence similarity and limit them to the reported 25%, we see identical Pearson Correlation coefficients for ThermoMPNN^47^ (0.34 versus 0.35) and a decreased correlation coefficient for RaSP (0.21 versus 0.30). This suggests that RaSP^21^ is more sensitive to the input data, with a bias towards the PDB structures Rosetta was optimized for.

We found a strong correlation between the datapoints in the S612 dataset with the CPA2 data. When further investigating the correlations of this to the CPA2 predictions, FoldX benefits the most from using the MD ensemble as input. Interestingly, the coarse grain ensemble generated with CABSflex^57,69^ is worse than the single structure, indicating that the movements of the backbone may exaggerate fluctuations creating states that are unrealistic and unstable, thus negatively affecting the change in stability predictions. The case of CPA2 is also a cautionary reminder, that navigating the growing field of experimental data and computational developments come with new challenges of overlap. Most recently Beltran et al.^74^ published a paper on site-saturation mutagenesis on hundreds of domains concurrently, where multiple proteins from the S669 dataset is included, which should make developers cautions when splitting their data for training and testing.

In conclusion, we are approaching an exciting time for stability predictors, where the expansion of experimental data may lead to model development with novel accuracy and speed. The integration of novel data into machine learning models, the improvement in computational efficiency, and the challenges surrounding benchmark data stress the dynamic nature of the field. To advance further, an effort to ensure that the models are trained on rigorously partitioned data that accounts for both sequence and fold similarity is needed, helping to address methodological biases. This might pave the way for more accurate and reliable models in computational biology. Additionally, it might be beneficial to apply some of the logic used in other bioinformatic fields^65,75–77^, where multiple methods are combined aiming to overcome the limitations of a single tool, or as applied in the MAVISp framework to rely on classification consensus between methods^49^. Finally, we highlight that the choice of the input structure is a critical parameter, and our results demonstrate that using molecular dynamics ensembles as input may enhance predictive performance^37^.

## Materials and Methods

### S4038 dataset & subsampling

The S4038 dataset^50^ were downloaded from the supplementary material of the paper. The subsampling protocol is based on the average value of samples from proteins, average occurrence of substituted amino acids and number of stabilizing mutations, where ΔΔG =< 0 kcal/mol is defined as stabilizing and ΔΔG > 0 kcal/mol is defined as destabilizing. We do not give any weights to specific species. We conduct this subsampling ten times to overcome bias of a single subsample set. For the average number of mutations identified in the ten rounds of under sampling we collect ten datasets of random sampling as controls.

### S669 dataset Curation, Structure Choice & Subsampling

We downloaded the S669 dataset curated by Fariselli’s group^54^. The S669 dataset comprises mutations from the ThermoMut Database (ThermoMutDB)^58^. The data contained in S669 are novel variants unlikely to have been used as training sets to develop methods to predict changes in ΔΔG upon mutation. The reasoning behind this is threefold: i) From the entire ThermoMutDB^58^, Fariselli’s group filtered away any mutation from three of the large and well-known datasets, included in ProTherm^60^, S2648^61^, and VariBench^62^. ii) From the remaining dataset, the proteins were excluded if they had more than 25 % sequence identity with the above-mentioned datasets. iii) Any inconsistent mutations were removed. Inconsistencies included multiple variants; the reported ΔΔG was in transition state kinetics, etc. These three steps left the 669 variants. Thus, the S669 dataset has been carefully manually curated. The aim is to study predicted structures, i.e. AlphaFold2 structures, since these increasingly lay the foundation for computational protein biology. Hence, we traced back each mutation to the ThermoMutDB for the uniprot accession number and searched the AlphaFoldDB^37,78^ for the predictive structure. Here we identified corresponding structures for each protein; we trimmed the AlphaFold model to remove disordered tails and linkers. Eight proteins were excluded, one due to inconsistencies with AlphaFold models (uniprot accession number P01837; 1 mutation) being a murine monoclonal antibody where the mutations refer to a specific PDB, and seven due to missing Alphafold structures (Uniprot Accession numbers: P11532; 1 mutation, P04517; 5 mutations, P12497; 1 mutation, P26747; 2 mutations, P03322; 4 mutations, P12823; 4 mutations, P69168; 27 mutations). We aligned all the used PDBs to the corresponding AlphaFold structure, annotating the listed mutation with its AlphaFold position and retained only mutations occurring at sites with an pLDDT score of or above 70, only including high and very high confidence regions. This left 615 mutations across 77 proteins, where 184 mutations across 31 proteins are human. Three of these mutations, on the Podospora anserina (Pleurage anserina) protein het-s (UniProt accession number: Q03689) feel outside the trimming region of the predictive runs leaving 612 mutations. While S612 overcomes the redundancy of benchmarking with the same or similar variants used for training or optimizing, it does not overcome the data imbalance concerning representation of amino acids, specific proteins, and imbalance of positive and negative cases^41^. Therefore, we balance the S612 dataset in relation to represented proteins, substituted amino acids and destabilizing mutations by down sampling the data. The down sampling follows the same procedure as S4038. Additionally, we checked the sequence similarity between the predictor ThermoMPNN’s training set and the proteins in S612. We identify 280 mutations across 19 proteins with a sequence similarity of 0.25 or higher, this includes UniProt accession numbers: P00149, P00441, Q9RA57, P37957, P18429, P0A780, Q97ZL0, O66523, P02654, P51161, Q9GZQ8, P0A3C7, Q5SIY4, P11961, Q53291, P05112, Q03689, P02417 and P0A9D2, which are excluded in the S332 reporting.

### Ensemble Generation

We generate two ensembles, one using CABS-flex^57,69^ and one using GROMACS^79^ molecular dynamics. The All-atom molecular dynamics simulation of CPA2 were performed for one microsecond using the CHARMM36m force field^80^. Following equilibration, the simulation was conducted in the canonical ensemble with explicit solvent and periodic boundary conditions. The generated ensemble underwent quality control assessment using Mol_Analysis^81^ and 25 frames extracted for stability predictions. The CABS-flex ensemble was generated using the CABS-flex 2.0 method and software within a snakemake pipeline available at https://github.com/ELELAB/MAVISp_CABSflex_pipeline. For CPA2 we restrained the secondary structure to avoid extreme conformations and a quality control step to evaluate the secondary structure content of the generated structures with respect to the starting one, using DSSP^82^and the SOV-refine score^83^. The method yielded 20 frames for analysis representing the 20 largest clusters.

### CPA2 dataset

The additional stability dataset of the Carboxypeptidase A2 activation domain (CPA2) is available in the maveDB^66,68^ - urn:mavedb:00000164-0-1 Published Jul 03, 2023 and part of the mega-scale dataset^2^. The data is obtained from a measure of the digestion of CPA2 protein variants with both trypsin and chymotripsin to measure the effect of these variants on folding. The score column used in this study represent the ΔG value, fitted using a baysian model, calculated using the coupled model from the counts across replicates and proteases. We aligned the sites to the AlphaFold model, and calculated the ΔΔG, ΔΔG = ΔGwt - ΔGmut, where the wild-type ΔG were established as an average of the six measurements reported in the dataset.

### Stability Predictors

The chosen stability predictors include the sequence based DDGun^11,84^, and the structure based DDGun 3D^11,84^ alongside FoldX in the MutateX implementation^56^, Rosetta in the RosettaDDGprediction implementation^63^, RaSP^21^ and ThermoMPNN^47^. The sequence based DDGun was developed as a baseline benchmarking tool and predicts the stability alteration because of mutation using a weighted linear combination of features^11^. These features include the Blosum substitution Matrix^70^ describing evolutionary likelihood, Skolnick’s statistical potential^71^ describing the difference in interaction energy in a sequence window of two residues and Kyte-Doolittle’s scale^72^ to describe the difference in hydrophobicity of the substitution. We use a stand-alone version and use the entire sequence of the protein as input. Additionally, for DDGun 3D, a fourth structural element is introduced^11^. This structural element is calculated with the Bastolla-Vendruscolo statistical potential^85^ describing the change in interaction energy within a sphere around the variant site with a radius of 5Å while the Cα position is unchanged. We apply the stand-alone versions for both. The MutateX implementation^56^ of FoldX^20^ provides a rigorous protocol to the FoldX methodology enforcing multiple rounds of calculations to find an average stability change upon mutation. FoldX predicts free energy changes of folding upon mutation by comparison of the wild-type and mutated structure, relying on an empirical free energy function including terms such as Van der Waals forces and electrostatic contributions. Notably the backbone of the protein remains stiff, and the motion of the mutated residue is limited to the rotamers. The RosettaDDGprediction^63^ implementation of Rosetta is, similarly to MutateX, a wrapper introducing standardized Rosetta protocols. The stability protocols aim to determine the stability alteration upon mutation as a change in gibbs free energy^17–19^. The protocols sample the structure in the cartesian space to generate a limited structural ensemble allowing some local backbone movement^17–19^. Rosetta also relies on an energy function^17–19^ with terms such as Lennard-Jones attractions and repulsions, Lazaridis-Karplus solvation energy, and coulombic electrostatic potential. We apply the rosetta ref 2015 protocol. RaSP^21^, the abbreviation for Rapid protein stability prediction, is a deep learning tool aiming to replicate the stability predictions of Rosetta. RaSP is a combination model utilizing supervised and self-supervised models, where the self-supervised step relies on a 3D convolutional neural network for a representation model which is in turn the input for the supervised model aiming to refine the output based on stability changes predicted by Rosetta with the ‘cartesian_ddg’ protocol^17–19^, why it is trained to replicate Rosetta. We apply a stand-alone version of RaSP adapted for application in the MAVISp framework^49^. Lastly, we apply ThermoMPNN^47^, is a message-passing neural network trained on a mix of the mega-scale data^2^ and FireprotDB^59^ where the FireprotDB have significant overlap with the S2648 dataset, containing 2648 single point mutations on 131 globular proteins^61^. The tool is built on the ProteinMPNN^86^ framework and we used the stand-alone version.

Each stability predictor estimates the stability change as ΔΔG in kcal/mol, and we invert the value if needed so a stabilizing mutation < 0 and a destabilizing mutation > 0.

### Prediction Evaluation

To evaluate the performance of the predictors we calculate the Pearson and Spearman correlation coefficients^87^ correlating the experimental and predicted values. The Pearson correlation measures the strength of an assumed linear relationship between the variables and is sensitive to outliers and non-linear relationships^88^. The Spearman correlation coefficient does not assume a linear relationship and is therefore less sensitive towards outliers^88^. We should assume a linear relationship, as the predictors aim to predict the experimental values, why a Pearson correlation is suitable, and is the primary reported correlation coefficient. However, the Pearson correlation coefficient aims to establish a linear relationship between normally distributed variables^88^, which stability data seldom have, due to the overrepresentation of destabilizing variants. Hence, Spearman is included in the supplementary materials. Additionally, we calculate Mean Absolute Error (MAE), thus measuring the average absolute difference between the experimental and predicted values. The MAE provides a straightforward measure of the average prediction error, which should be as low as possible^89^.

## Data Availability

Code to replicate figures and novel data for this publication is available at: github.com/ELELAB/Stability_Predictors_Benchmark

## Author Contributions

*Conceptualization*: EP, KD; *Data Curation*: KD, MU, PSI. *Formal Analysis*: KD. *Funding Acquisition:* EP, *Investigation*: KD. *Methodology*: KD, MT, EP. *Supervision*: EP, MT. *Visualization*: KD. *Writing:* KD. *Review and Editing*: PSI, MU, MT, EP.

