## supplementary_figures1_2 for "In-Silico Stability Predictors: Investigation of Performance Towards balanced Experimental Data"

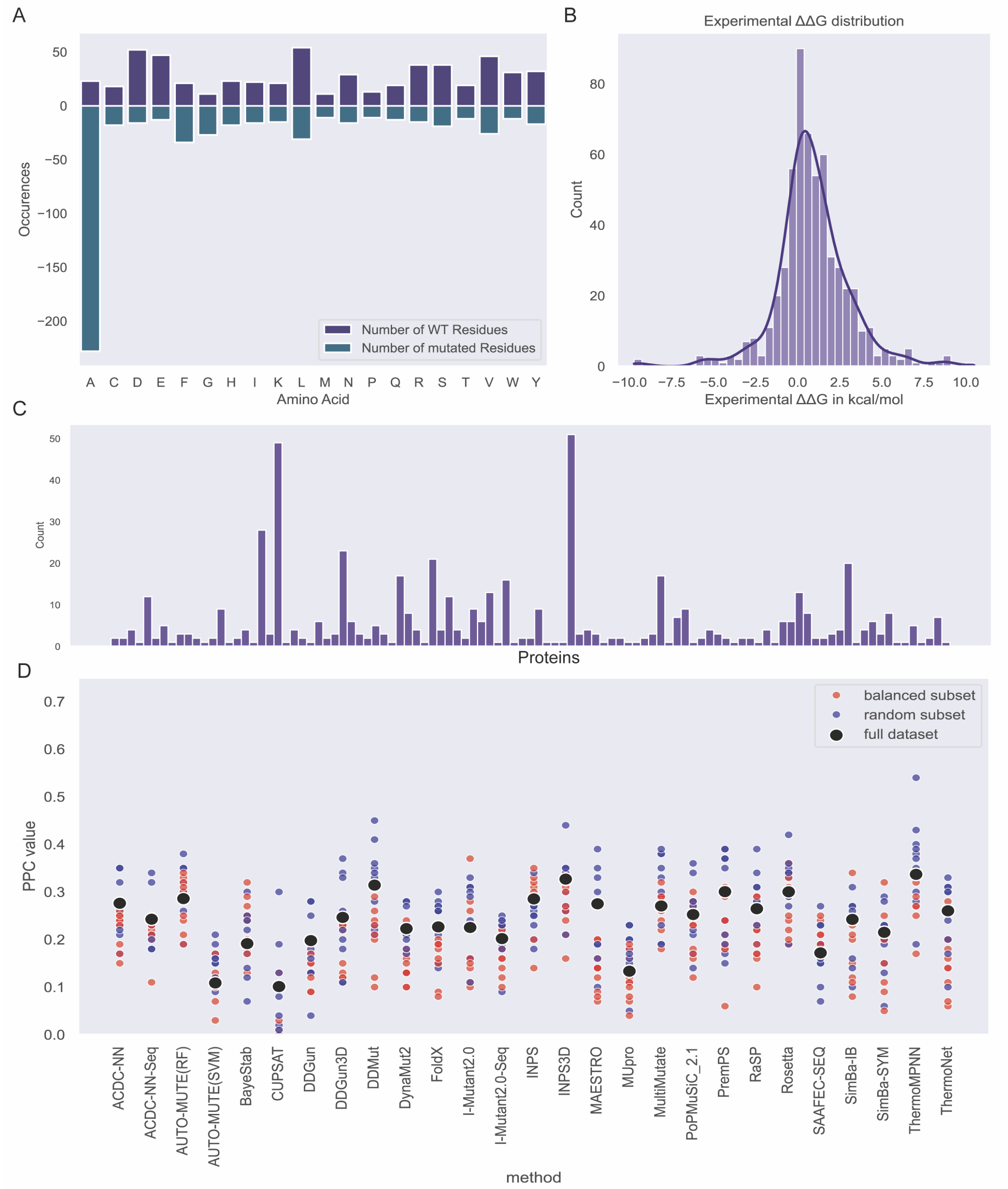


**Figure S1:** Distributions of S4038 excluding annotated mutations existing in specific training sets, collectively 568 mutations (A) Amino Acid count. The plot illustrates the number of sites mutated from a wild-type (WT) amino acid on residues of the sites in the dataset in purple, positive y-axis, and number of sites mutated to an amino acid, in teal, negative y-axis. (B) The histogram illustrates the distribution of the experimental ΔΔG values, showing a higher number of destabilizing mutations (ΔΔG > 0) than stabilizing mutations (ΔΔG < 0) (C) The bar chart illustrates each of the proteins on the x-axis and the number of mutations reported in each protein. (D) A scatterplot of the Pearson Correlation Coefficient (PCC) of all the 568 mutations (black), and the balanced subsets (red) and random subsets (blue). For many of the benchmarked tools (x-axis), there is not as clear of a separation between the balanced subsets and the random subsets, while there still is a definite tendency of the balanced subsets tend to have lower PCCs towards the experimental data.

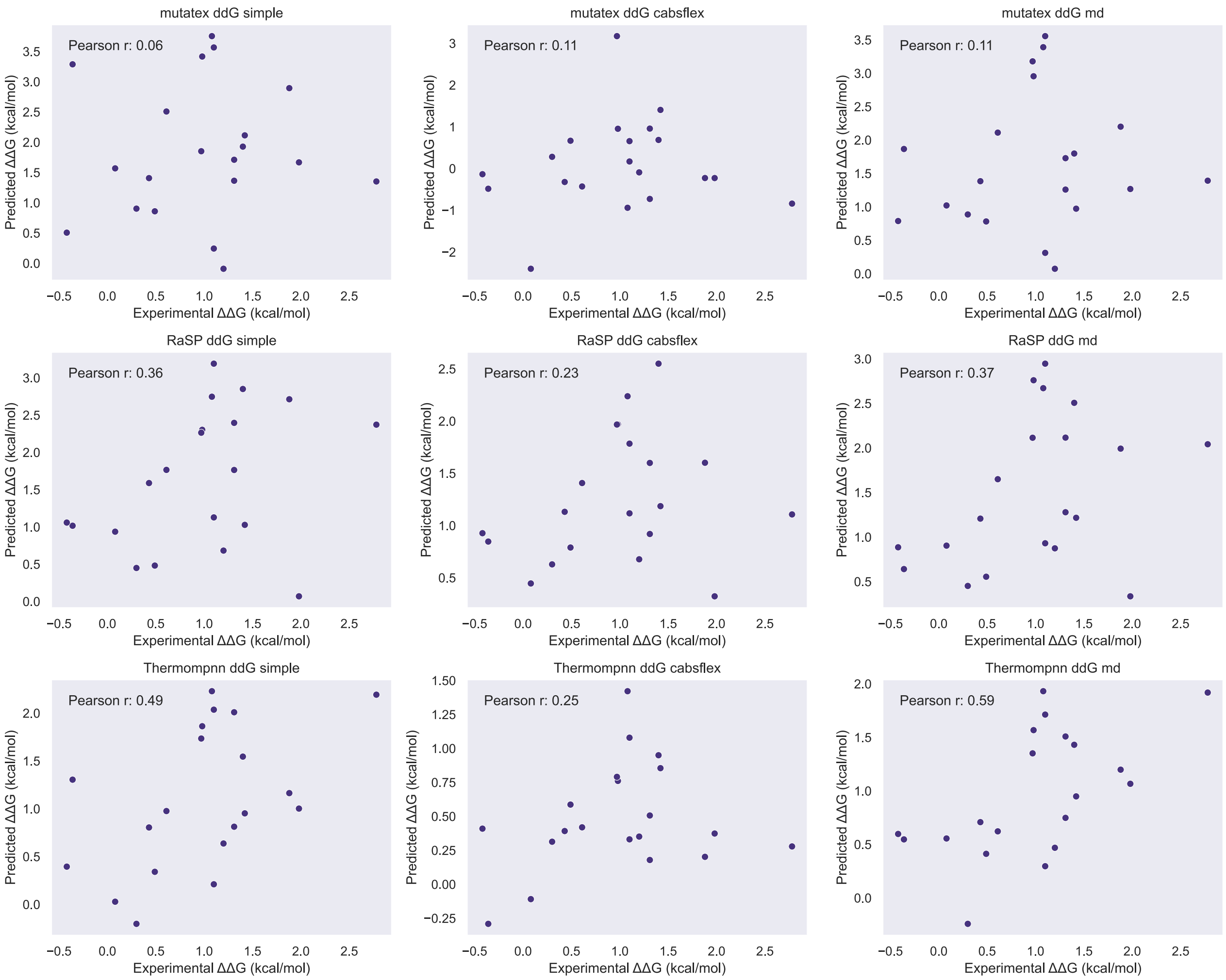


**Figure S2:** Scatterplots of the ΔΔG values from the S612 dataset and the predicted ΔΔG values on three different inputs, the simple (static) structure from AlphaFold, a CABSflex ensemble and a Molecular Dynamics ensemble.
